# Purification of functional mouse skeletal muscle mitochondria using Percoll density gradient centrifugation

**DOI:** 10.1101/2023.07.11.548594

**Authors:** Rea Victoria P. Anunciado-Koza, Anyonya R. Guntur, Calvin P. Vary, Carlos A. Gartner, Madeleine Nowak, Robert A. Koza

**Affiliations:** MaineHealth Institute for Research, 81 Research Drive, Scarborough, ME, United States of America; Graduate School of Biomedical Sciences and Engineering, University of Maine, Orono, ME, United States of America; Department of Medicine, Tufts University School of Medicine, Boston, MA, United States of America; Pennington Biomedical Research Center, 6400 Perkins Road, Baton Rouge, LA, United States of America

**Keywords:** Percoll purification, density gradient, mitochondria, bioenergetics, skeletal muscle

## Abstract

**Objective:** Our goal was to isolate purified mitochondria from mouse skeletal muscle using a Percoll density gradient and to assess bioenergetic function and purity via Seahorse Extracellular Flux (XF) Analyses and mass spectrometry.

**Results:** Mitochondria isolated from murine quadriceps femoris skeletal muscle using a Percoll density gradient method allowed for minimally contaminated preparations with time from tissue harvest to mitochondrial isolation and quantification in about 3-4 hours. Percoll purification from 100-200 mg fresh tissue yielded ∼200-400 ug protein. Mitochondrial bioenergetics evaluated using the Seahorse XFe96 analyzer, a high-throughput respirometry platform, showed optimum mitochondrial input at 500 ng with respiratory control ratio ranging from 3.9-7.1 using various substrates demonstrating a high degree of functionality. Furthermore, proteomic analysis of Percoll-enriched mitochondria isolated from skeletal muscle using this method showed significant enrichment of mitochondrial proteins indicating high sample purity. This study established a methodology that ensures sufficient high quality mitochondria for downstream analyses such as mitochondrial bioenergetics and proteomics.

## Introduction

In humans, the skeletal muscle accounts for 40% of total body mass and plays important roles in locomotion and metabolic function (1). In earlier studies using uncoupling protein 1 (UCP1)-deficient mice, we focused on the potential involvement of skeletal muscle in UCP1-independent thermogenesis (2). Mitochondria, which form a three-dimensional network throughout the cell, are one of three important cellular organelles in the muscle fiber along with the transverse tubular system and sarcoplasmic reticulum (1). Complex interconnected mitochondria lead to higher oxidative capacity while isolated ellipsoidal mitochondria are associated with less efficient mitochondrial respiration (3). Differential centrifugation is one of the most commonly used methods for skeletal muscle isolation of mitochondria for metabolic studies (4). Unlike differential centrifugation where most skeletal mitochondrial preparations are contaminated with other organelles (lysosomes, peroxisomes, Golgi membranes, sarcoplasmic reticulum), density-gradient centrifugation with Percoll leads to highly purified preparations while maintaining respiratory activity (4, 5). The purity of the mitochondrial preparation can influence the results of down-stream analyses. In this study, our goal was to obtain a high yield, high purity mitochondrial preparation whilst maintaining viability by evaluating the respiratory control ratio (RCR) using a Seahorse XFe96 Extracellular Flux Analyzer and concurrently performing proteomic analyses to demonstrate purity.

## Main Text

### Materials and Methods

#### Isolation and Purification of Mitochondria

Animal experiments and methods were approved by the MaineHealth Institute for Research Institutional Animal Care and Use Committee. All buffers and labware were pre-chilled on ice. Chemicals were obtained from Sigma-Aldrich (St. Louis, MO, USA) unless otherwise indicated. Reagents for isolation of mitochondria are listed in Additional File 1: Table S1.

Eight to ten week old mice were euthanized by cervical dislocation and dipped in 70% ethanol to minimize hair contamination in samples. Exposure to anesthesia prior to sample collection has been reported to impact mitochondrial respiration (6, 7) and therefore was not used. Skeletal muscle mitochondria was isolated by differential centrifugation (8) and Percoll gradient purified essentially as described (9) with some modifications. Quadriceps femoris skeletal muscles were collected and contaminating connective and adipose tissue were removed. Samples were immediately placed in a 50 ml polypropylene tube containing 20 ml of ice cold 1X DPBS (Lonza, Basel, Switzerland). Tissues were then transferred to a 50 ml beaker containing ice-cold 10 ml 1X DPBS, swirled briefly to remove blood, and decanted with the excess liquid blotted with a Kimwipe. Tissues were minced using scissors in 0.5 ml IM buffer containing Nagarse (0.6 mg/ml; Sigma-Aldrich, P8038) for 2 minutes. Another 0.5 ml of IM buffer/Nagarse was added, and the minced tissues were incubated at RT for 5 minutes. Using a pre-cut 1 ml pipette tip, digested tissues were transferred to a 7 ml Kimble Dounce tissue grinder. After the addition of 0.5 ml IM, tissues were homogenized using a loose fitting pestle for 10 strokes. Another 1 ml of IM buffer was added to each homogenate for additional 1-3 strokes. The homogenates were transferred to a 15 ml polypropylene tube. The homogenizers were rinsed with ice-cold 2.5 ml IM buffer/0.5% fatty acid-free BSA and combined with initial homogenates. Samples were centrifuged at 1000g at 4°C for 5 minutes. The supernatants were transferred to an ice-cold 10 ml polycarbonate centrifuge tube while avoiding the flocculent layer on top of the pellet. An additional 4 ml of IM buffer was added to each pellet and samples were briefly vortexed to resuspend the pellets and then centrifuged at 1000g at 4°C for 5 min. The supernatants were collected while avoiding the flocculent layer above the pellet, pooled with the previously collected supernatants, and then centrifuged at 21,000g at 4°C for 10 min. At this time, discontinuous Percoll gradients were prepared by pipetting 3.7 ml of 24% Percoll in 1X IM buffer (pH 7.4) onto the bottom of 10 ml polycarbonate centrifuge tubes followed by carefully pipetting of 1.5 ml of 40% Percoll in 1X IM buffer (pH 7.4) under the 24% Percoll layer maintaining a sharp interface between layers. The supernatants from the centrifuged (21,000g) samples were removed and discarded, and pellets were resuspended in 2 ml of 15% Percoll in 1X IM (pH 7.4) by gentle pipetting then filtered through a pre-wetted (0.5 ml 15% Percoll) 70 μm nylon filter. The centrifuge tubes were washed with an additional 1 ml of 15% Percoll that was also passed through the filter. The 3.5 ml of 15% Percoll filtrates containing mitochondria were carefully layered on top of the 10 ml polycarbonate centrifuge tubes containing the discontinuous Percoll gradient while maintaining a sharp interface between the 15% and 24% Percoll layers. Samples were then centrifuged at 30,750g at 4°C for 10 min using slow acceleration and no deceleration (brake). After centrifugation, around three bands were evident. The upper layers were removed and discarded and the enriched mitochondrial layer between the 24% and 40% Percoll interface was collected and placed into a 10 ml polycarbonate centrifuge tube. Following the addition of 6 ml of IM buffer, the enriched mitochondrial fraction was centrifuged at 16,750g at 4°C for 10 min using maximum acceleration and slow deceleration. The supernatant was carefully removed and discarded while being careful not to disturb the loose mitochondrial pellet on the bottom of the tube. The pellet was resuspended in 1 ml of IM buffer containing 0.5% fatty acid-free BSA and then 4 ml of IM buffer containing 0.1% fatty acid-free BSA, mixed, and centrifuged at 7000g at 4°C for 10 min. Supernatants were decanted and discarded and residual solution from the wall of the centrifuge tube was removed. Mitochondrial pellets were resuspended in 0.75 ml IM buffer, transferred to 1.5 ml microfuge, and the centrifuge tube was washed with an additional 0.75 ml IM and combined with the initial suspension. Samples were centrifuged in a microfuge at 10,000g at 4°C for 10 min. Supernatants were discarded and pellets gently resuspended in 100 μl of 1X MAS buffer without BSA. Aliquots (e.g. 5 μl) were removed for protein quantification using a Qubit Protein Assay Kit (Life Technologies, Carlsbad, CA, USA), and additional aliquots were saved for downstream applications requiring the absence of BSA (e.g. proteomics, western blotting etc.). An equal volume of 1X MAS buffer supplemented with 1% fatty acid-free BSA was added to the remaining fractions of enriched mitochondria and mitochondrial suspensions were maintained on ice prior to being used for measurements of mitochondrial respiration.

### Measurements of mitochondrial respiration using extracellular flux (XFe96) assay

In order to determine functionality of the isolated mitochondria, respirometry was performed immediately after isolation using the Seahorse XFe96 platform (Agilent Technologies) as previously described with slight modifications (8, 10). The substrates, inhibitors and assay solutions used are listed in Additional File 2: Table S2. For detailed assay procedures, see Additional File 3: Methods.

### Mitochondrial proteomics

Briefly, tryptic peptides from Percoll gradient-enriched mitochondrial protein were run on a Sciex TripleTOF 5600 mass spectrometer connected to a Dionex Ultimate 3000 (RSLCnano) chromatography system. The mass spectrometer was operated using data dependent acquisition (DDA) to create an ion library. A Sequential window acquisition of all theoretical spectra (SWATH) was implemented for relative quantitation, as previously described (11-13). Spectral alignment and targeted extraction of SWATH data were performed with the SWATH Processing Micro App in PeakView (Version 2.2.0, Sciex) using the reference DDA ion library. The data was imported into MarkerView (Version 1.2.1, Sciex), where data was ratio-normalized, groups compared by principle component analysis, and compared for significance using the Fisher’s modification of Student’s T-Test (14). Detailed methodology for mitochondrial proteomics are described in Additional File 3: Methods.

### Statistical analysis

Data are expressed as mean ± SEM or mean ± SD using MS Excel 2016 and Graphpad Prism 9.4.

## Results and discussion

The general scheme for the isolation of mouse skeletal muscle mitochondria is presented in Fig. 1. We collected mouse quadriceps femoris muscle from both legs and processed by differential centrifugation to obtain the crude mitochondrial fraction that was further purified by Percoll density gradient to remove contaminating organelles. Time from tissue harvest to mitochondrial isolation and quantification is ∼3-4 hours. Percoll purification is compatible with fresh tissues collected from each mouse yielding 200-400 μg protein from 100-200 mg tissue. The high mitochondrial protein yield from fresh samples ensures sufficient material for downstream analysis such as mitochondrial bioenergetics and proteomics. Metabolic studies of mitochondria isolated from various tissues are often prepared by differential centrifugation, avoiding use of density media (sucrose, Percoll, Nycodenz, Iodixanol, Ficoll) which prolongs the preparation time (4). However, mitochondria prepared by differential centrifugation are often contaminated to some extent with organelles such as lysosomes, peroxisomes, Golgi membranes and endoplasmic reticulum (4). In the interest of obtaining an enriched mitochondrial preparation, while maintaining functionality, we used Percoll gradient purification. Percoll, a colloidal suspension of silica particles coated with polyvinylpyrrolidone, is nontoxic, chemically inert and has negligible osmolarity (5, 15). Percoll is applicable to any cells or organelles in suspension for which differences in size or density exist (15). It has been used in isolation of mitochondria from various tissues such as brain, heart, skeletal muscle and liver (4, 5, 9). Moreover, the Percoll density gradient method does not require ultracentrifugation and is compatible with more commonly used medium or high-speed centrifuges with fixed angle rotors (9).

**Figure 1.**
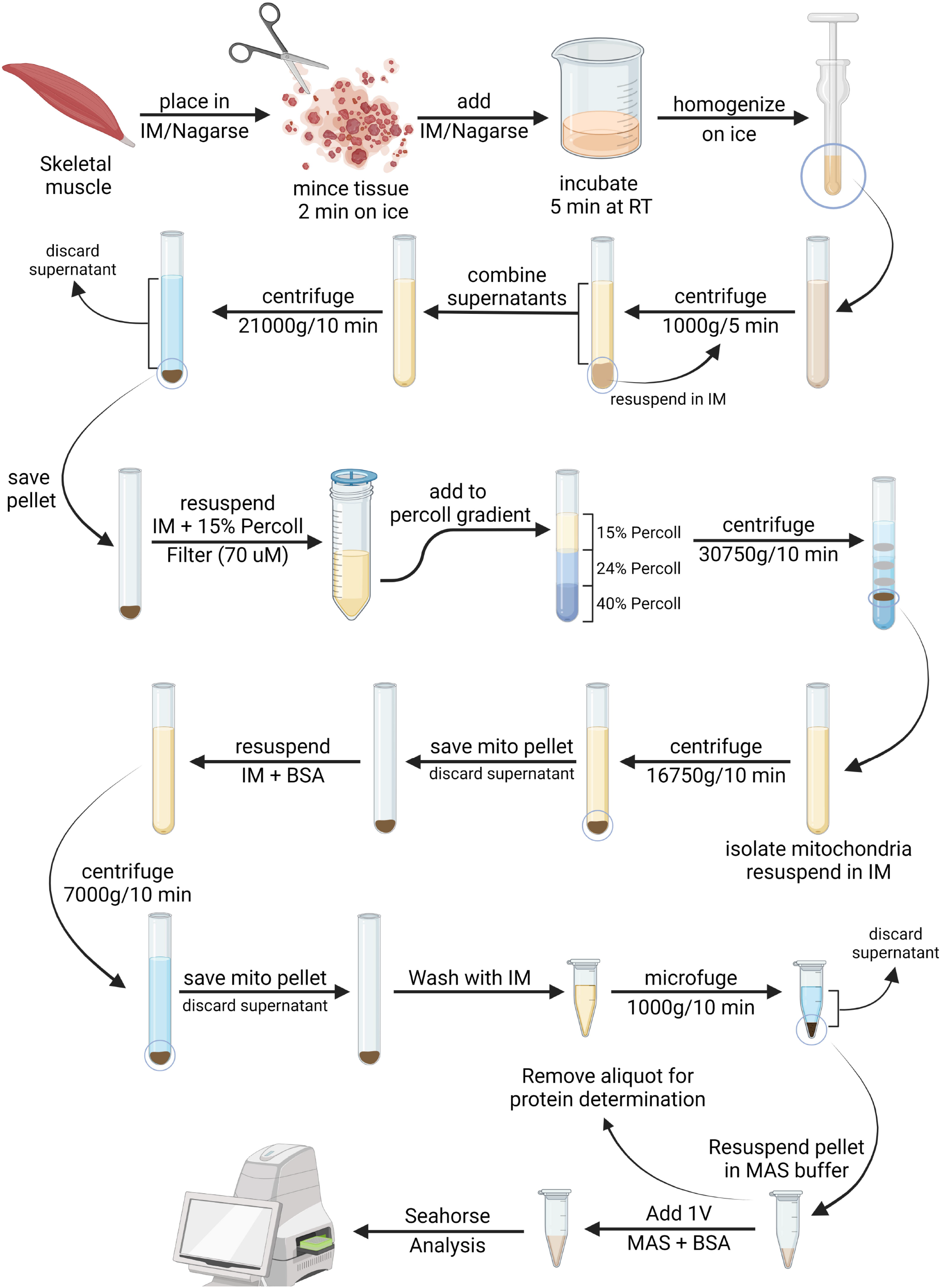
Schematic representation of enrichment procedure for mitochondria from mouse skeletal muscle. Illustration created using icons from BioRender (https://biorender.com/).

Immediately after mitochondrial isolation and quantification, we performed respirometry on the samples using XFe96 format. To determine functionality of the isolated mitochondria based on analysis of respiratory rates and coupling, different amounts of mitochondrial protein were loaded in each well (1000, 500, 250 and 125 ng) in the presence of succinate/rotenone as substrate (Fig. 2a). Basal respiration (State 2) is linear to the mitochondrial input; however, optimal OCR occurred with a 500 ng protein input. In addition, overload of mitochondrial input at 1000 ng was determined by examining the O_2_ tension (in mmHg) with almost complete depletion of O_2_ in the microchamber (Fig. 2b). RCR, a gauge for mitochondrial coupling, ranged from 3.6-4.1 (Fig. 2a). We also performed coupling assays using different substrates/<u>additives</u>) of the electron transport chain (Complex I: pyruvate/<u>malate</u>, glutamate/<u>malate</u>, palmitoly L-carnitine/<u>malate</u>; Complex II: succinate/<u>rotenone</u>) (Fig. 2c, 2d) which showed RCR’s ranging from 3.9-7.1 indicating intact mitochondria following Percoll gradient enrichment. Owing to the high quality of mitochondria obtained from this method, coupling assays to interrogate the different mitochondrial complexes can be performed simultaneously with very small amounts of material.

**Figure 2.**
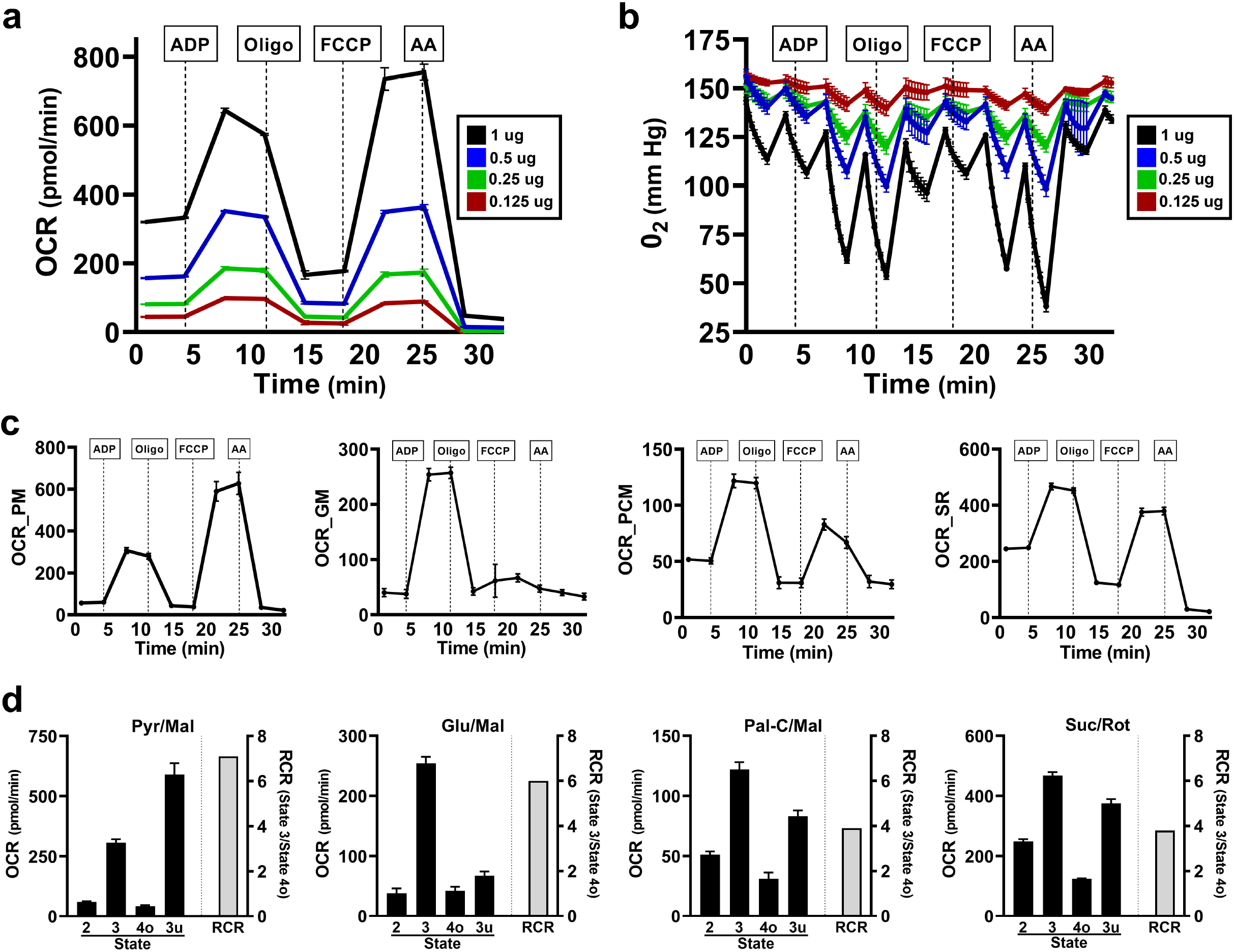
Bioenergetic profile of coupling assays in isolated mitochondria. (a) Determination of optimal mitochondrial input per well (1000, 500, 250 125 ng) in the presence of succinate/rotenone. (b) Absolute O_2_ tension (in mm Hg) in microchamber with different mitochondrial inputs. (c) Coupled mitochondrial respiration assays using 10 mM pyruvate/2 mm malate (PM), 10 mM glutamate/10 mM malate (GM), 40 uM palmitoyl L-carnitine/1 mM malate (PCM) and 10 mM succinate/2 uM rotenone (SR) as substrates with 600 ng mitochondrial input per well. (d) OCR for States 2, 3 4o and 3u and RCR for each of the substrates in (c). For Figures (a-c) data are expressed as pmol/min ± SEM or mm Hg ± SEM and in (d), as pmol/min ± SEM or the ratio of State 3/State 4o (RCR).

Analyses of an aliquot of a mitochondrial preparation from skeletal muscle of a C57BL/6J mouse by peptide quantitative SWATH mass spectrometry identified 409 proteins (Additional File 4: Proteomics) with approximately 81% (n=333) showing association with mitochondria. Importantly 99% of the top 100 proteins, based on the number of peptide hits, identified with mitochondria. Of the top 250 proteins, 93.6% showed association with mitochondria with most (55.2%) compartmentalized within the mitochondrial matrix (Fig. 3). Other major categories included general mitochondrion (20.8%), mitochondrial inner membrane (13.6%), mitochondrial inner membrane space (2.0%) and mitochondrial outer membrane (2.0%). Cytosolic, cytoplasm, keratin filament and plasma membrane proteins made up the bulk (5.2%) of the remaining 6.4% of the non-mitochondrial proteins.

**Figure 3.**
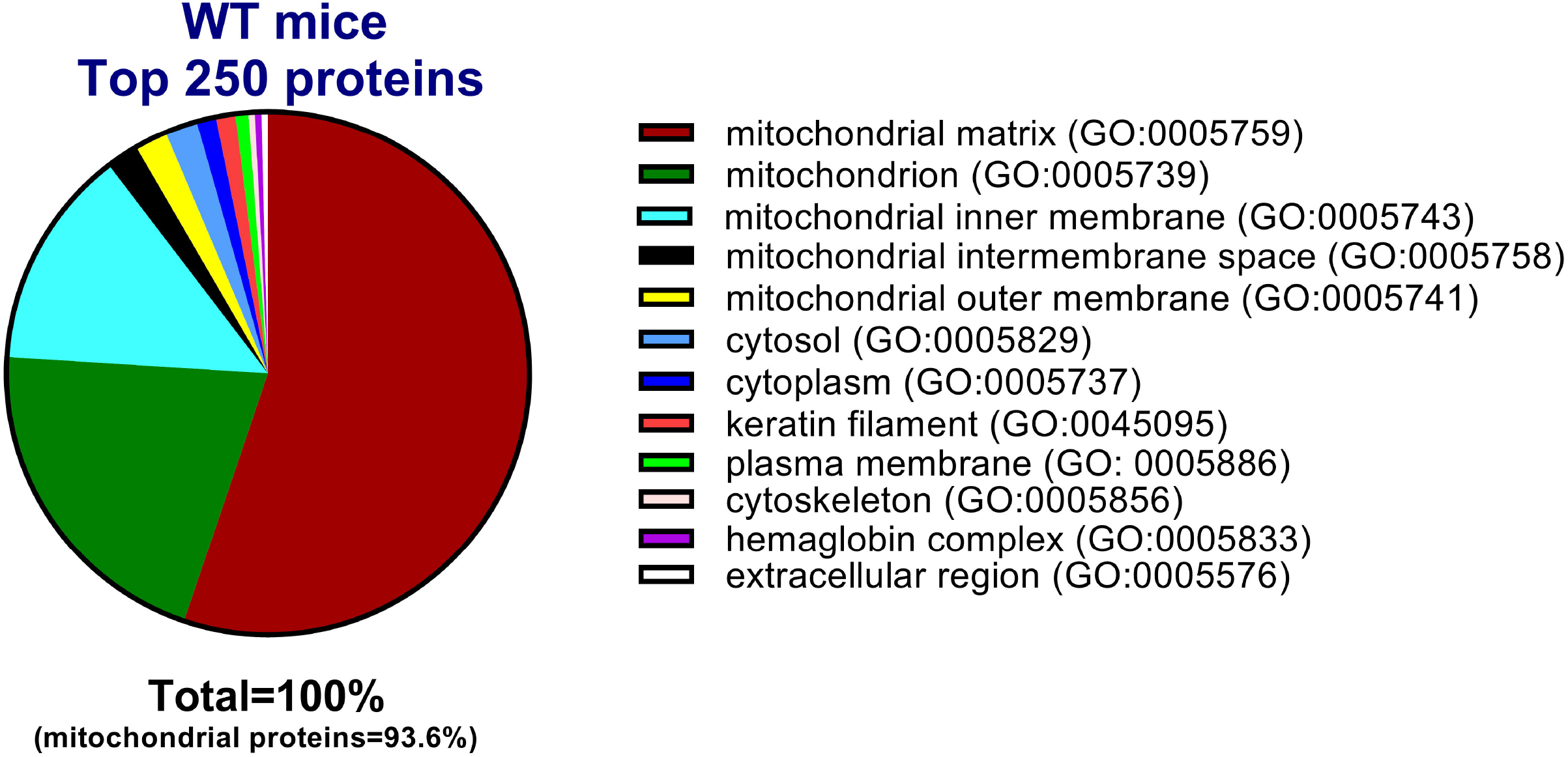
SWATH-MS identification of proteins in enriched mitochondrial preparations from skeletal muscle. A pie chart represents gene ontology (GO) categories of proteins identified in Percoll gradient-enriched mitochondria isolated from quadriceps femoris skeletal muscle of a C57BL/6J mouse.

In this study, we optimized a protocol to maximize the use of mitochondria isolated from 100-200 mg of mouse skeletal muscle for a microplate-based high-throughput respirometric assay. We also demonstrated a high degree of functionality with minimal contamination of other subcellular fractions as shown by proteomics analysis.

## Limitations

A limitation of this study is isolation of the total mitochondrial population in a mixed type tissue (quadriceps femoris) composed of type 1 (red) and type 2 (white) skeletal muscle fibers. Moreover, subsarcolemmal and intermyofibrillar mitochondrial subpopulations of skeletal muscle were not distinguished.

## Declarations

### Ethics approval and consent to participate

All animal experiments and methods were reviewed and approved by the MaineHealth Institute for Research (MHIR) Institutional Animal Care and Use Committee (IACUC). The study is in accordance with National Institutes of Health guidelines for the care and use of laboratory animals.

### Consent for publication

Not applicable. This study does not contain human studies or data.

## Availability of data and materials

All relevant data are contained within the manuscript and/or supporting information files.

## Authors Contributions

R.V.P.A.K., R.A.K and A.R.G. designed the study and performed the experiments. C.V. and C.A.G. performed peptide quantitative SWATH-MS and data analyses. R.V.P.A.K. and R.A.K. wrote the manuscript text. M.N. and R.A.K. designed and created figures 1-3. All authors reviewed the manuscript.

## Acknowledgements

R.A.K. and G.A.R. were supported by NIH/NIGMS P20GM121301 and the MaineHealth Institute for Research (MHIR). The Physiology and the Proteomics and Lipidomics Cores at MHIR were supported by NIH/NIGMS P20GM121301. The content is solely the responsibility of the authors and does not necessarily represent the official views of the National Institutes of Health.

