## Additional File 1 for "Purification of functional mouse skeletal muscle mitochondria using Percoll density gradient centrifugation"

**Additional File 1: Table S1. Reagents for mitochondrial isolation**

| **Compound** | **Reagent** | **Source, Identifier** |
| --- | --- | --- |
| Chappel-Perry (CP) Buffer I | potassium chloride (100 mM) | Sigma-Aldrich, P9541 |
|  | MOPS (50 mM) | Sigma-Aldrich, M1254 |
|  | EDTA (1 mM) | Sigma-Aldrich, ED2SS |
|  | magnesium sulfate (5 mM) | Sigma-Aldrich, M2643 |
| Isolation Medium (IM) | sucrose (225 mM) | Sigma-Aldrich, S0389 |
|  | mannitol (75 mM) | Sigma-Aldrich, M4125 |
|  | EGTA (1 mM) | Sigma-Aldrich, E3889 |
|  | HEPES (5 mM) | Sigma-Aldrich, H3375 |
| Isolation Medium (IM) + BSA | fatty acid-free BSA (0.5% and 0.1%) | Sigma-Aldrich, A7030 |
| 1X IM w/ 15%, 24% and 40% Percoll | Percoll^®^ density gradient media | Cytiva, 17089102 |
