## Additional File 2 for "Purification of functional mouse skeletal muscle mitochondria using Percoll density gradient centrifugation"

**Additional File 2: Table S2. Reagents for Seahorse extracellular flux (XF) assay**

| **Compound** | **Reagent** | **Source, Identifier** |
| --- | --- | --- |
| Substrates and Inhibitors | succinic acid | Sigma-Aldrich, S3674 |
|  | rotenone | Sigma-Aldrich, R8875 |
|  | sodium pyruvate | Sigma-Aldrich, P8574 |
|  | L-malic acid | Sigma-Aldrich, 02288 |
|  | glutamic acid | Sigma-Aldrich, G1251 |
|  | ADP | Sigma-Aldrich, A2754 |
|  | oligomycin | Sigma-Aldrich, O4876 |
|  | FCCP | Sigma-Aldrich, C2920 |
|  | antimycin A | Sigma-Aldrich, A8674 |
|  | palmitoyl-L-carnitine chloride | Sigma-Aldrich, P1645 |
|  | sodium L-ascorbate | Sigma-Aldrich, A4034 |
|  | TMPD | Sigma-Aldrich, T3134 |
| Mitochondrial assay solution (MAS) | sucrose | Sigma-Aldrich, S0389 |
|  | mannitol | Sigma-Aldrich, M4125 |
|  | magnesium chloride | Sigma-Aldrich, M9272 |
|  | HEPES | Sigma-Aldrich, H3375 |
|  | potassium phosphate monobasic | Sigma-Aldrich, P5655 |
|  | fatty acid-free BSA | Sigma-Aldrich, A7030 |
