## Additional File 3 for "Purification of functional mouse skeletal muscle mitochondria using Percoll density gradient centrifugation"

**Additional File 3: Methods**

**Measurements of mitochondrial respiration using extracellular flux (XFe96) assay**

On the day prior to the assay, the XF cartridge was hydrated per manufacturer’s instructions using 200 ul of sterile tissue culture grade water (Lonza) at 37°C in a non-CO_2_ incubator. On the day of the assay, water was discarded and replaced with 200 ul pre-warmed XF calibrant (Agilent Technologies, Santa Clara, CA, USA) one hour prior to loading the injection ports of the cartridge. The hydrated cartridge was kept at 37°C in a non-CO_2_ incubator until calibration.

Reagents used for XF assay are listed in Supplemental File 2. All stocks were maintained at -20°C and working solutions were prepared fresh on the day of assay. Mitochondrial assay solution (MAS, 1X: 70 mM sucrose, 220 mM mannitol, 10 mM KH_2_PO_4_, 5 mM MgCl_2_, 2 mM HEPES, 1 mM EGTA and 0.2% fatty acid-free BSA) was prepared for dilution of substrates. BSA-free 2X MAS was prepared for dilution of injectables. Stocks (0.5 M) of succinate, malate, glutamate and ADP (80 mM) were prepared in water, and adjusted to pH 7.4 with potassium hydroxide. Palmitoyl L-carnitine chloride stock (10 mM) was prepared in 95% ethanol. Stocks of FCCP (10 mM), rotenone (2 mM), oligomycin (6 mM), and antimycin A (7.6 mM) were prepared in DMSO. Pyruvate was prepared fresh on day of assay. All substrates and injectables were adjusted to pH 7.4 at 37°C on the day of assay.

Respirometry in isolated mitochondrial samples were performed immediately after isolation. Isolated mitochondria were resuspended in IX MAS with 0.5% fatty acid-free BSA. The following substrates/additives were used for coupling assay and were prepared in 1X MAS/fatty-acid free BSA (final concentrations): pyruvate (10 mM)/malate (2 mM), glutamate (10 mM)/malate (10 mM), palmitoyl L-carnitine (40 µM)/malate (1 mM) and succinate (10 mM)/rotenone (2 µM). Compounds for injection were prepared in 1X MAS at 10X the final concentration required for the assay. The sensor cartridge were loaded with the following: 20 µl ADP (final concentration, 4 mM) in port A, 22 ul oligomycin (final concentration, 2 µM) in port B, 25 µl FCCP (final concentration, 4 µM) in port C and 28 µl antimycin A (final concentration, 2 µM). The cartridge was placed in the machine for sensor calibration. During sensor calibration, the isolated mitochondria were resuspended in the different substrates/additives and loaded in the assay plate. The assay plate was centrifuged at 2000g at 4°C for 20 minutes to ensure attachment of the mitochondria to the bottom of the plate. After completion of spin, pre-warmed (37°C) substrates/additives were carefully added to each well at 180 µl volume. Oxygen consumption rate of the isolated mitochondria was measured with the following protocol: 30s mix, 30s wait and 2 min measure cycles with 2 cycles after each injection. All respiratory measurements were performed on the XFe96 platform (Agilent Technologies). We assessed State 2 (basal respiration), State 3 (ADP-stimulated respiration), State 4o (oligomycin-induced respiration due to proton leak) and State 3u (maximal uncoupler-stimulated respiration) in a coupling assay using various substrates. Respiratory control ratio (RCR), an index of mitochondrial coupling was calculated by the ratio of State 3 and State 4o respiration.

**Mitochondrial proteomics**

**Trypsin digestion**

Approximately 30-50 µg of each Percoll gradient-enriched mitochondria preparation were re-suspended in 400 µl of 8M Urea/50 mM Tris-HCL pH 8 with protease inhibitors (Roche, Mannheim, Germany) then sonicated (Branson Sonifier 250, Branson Ultrasonics, Brookfield, CT, USA) for 3x10 seconds) and rested on ice. Each lysate was reduced for 30 minutes with 8 mM dithiothreitol and alkylated for 15 minutes with 20 mM iodoacetamide at 30°C. The 8M urea solution was diluted with 300 µl of 50 mM Tris-HCL pH 8, and samples were digested overnight with 20 µg sequencing grade trypsin (Trypsin Protease, MS Grade, Promega, Madison, WI, USA) (Optimal Ratio of Trypsin to Substrate 1:20 – 1:100). From each sample, 100 µl of the tryptic digest was normalized to approximately 30 µg by purification over C18 Spin Columns. (ThermoFisher Scientific, Rockford, IL, USA).

**Mass Spectrometry**

Tryptic peptides were re-suspended in 0.1% formic acid in 0.02% acetonitrile in LC-MS grade water. Tryptic digests were run on a Sciex TripleTOF 5600 mass spectrometer connected to a Dionex Ultimate 3000 (RSLCnano) chromatography system. Each sample was loaded onto an analytical reverse-phase C18 nanocolumn (40 cm length, 75 μm ID) packed with Reprocil Pur C18,1.9 μm and resolved by an increasing acetonitrile gradient over 100 min at a flow rate of 220 nl/min. The mass spectrometer was operated using data dependent acquisition (DDA) to create an ion library. A Sequential window acquisition of all theoretical spectra (SWATH) was implemented for relative quantitation, as previously described (1-3). For both DDA and SWATH workflows, the mass spectrometer used an ion spray voltage floating (ISVF) of 2400 V, curtain gas (CUR) 25 PSI, interface heater temperature (IHT) 150 °C, ion source gas 1 of 6 PSI and a declustering potential (DP) 100 V.

All data acquired in DDA mode used a high resolution MS scan from 350-1500 m/z to select the 50 most intense ions prior to MS/MS analysis using collision induced dissociation (CID). Charge states 2-5 were selected with an exclusion time of 15 seconds and an accumulation time of 250ms for TOF MS, and 50ms for TOF MS/MS, cycle time 2.8 seconds.

For SWATH acquisition, a TOF MS scan with an accumulation time of 96ms was followed by 100 variable-width scan windows from 350 to 1500m/z. Accumulation time was 89.9ms, cycle time 9.1 seconds. Identical chromatography parameters were used for SWATH and DDA analysis.

**Nano-Scale Liquid Chromatography**

During the first 0-9 minutes of chromatography, the sample was loaded onto a C18 trap column (Thermo Scientific) at a flow rate of 3 µl/min. At 9 minutes, elution began using a 1 minute period of buffer B increase from 0-1% at a flow rate of 220 nl/min. From 1-90 minutes, buffer B increases from 10-35%; from 91-95 minutes, buffer B increased from 35-95%, from 95-98 minutes, buffer B remained at 95%, and from 98-100 minutes, buffer B decreased from 95-2%. Buffer A: 98% water, 1.9% acetonitrile, 0.1% formic acid. Buffer B: 80% acetonitrile, 19.9% water, 0.1% formic acid. All reagents used were LCMS grade.

**Data Analysis**

For the creation of a peptide ion library, DDA data were searched against the reviewed mouse UniProt database (37,201 proteins) using ProteinPilot software (Version 5.0.2, Sciex) with the Paragon algorithm. Peptides used for protein identification met a minimum confidence threshold of greater than 99%.

Spectral alignment and targeted extraction of SWATH data were performed with the SWATH Processing Micro App in PeakView (Version 2.2.0, Sciex) using the reference DDA ion library. The data was imported into MarkerView (Version 1.2.1, Sciex), where data was ratio-normalized, groups compared by principle component analysis, and compared for significance using the Fisher’s modification of Student’s T-Test (4).
